# MICROPLASTICS IN THE MOUNT TERMINILLO SNOW’S (RIETI, ITALY)

**DOI:** 10.1101/2024.11.15.623744

**Authors:** Loris Pietrelli, Patrizia Menegoni, Ines Millesimi, Maria Sighicelli, Marina Baldi, Bartolomeo Schirone

## Abstract

Microplastics (MPs) concentrations in atmospheric fallout is a growing environmental concern, this paper reports data regarding the MPs pollution determined using snow samples collected in wintertime and in 6 different areas on Mount Terminillo (2216 m a.s.l.) a typical Apennines massif located north of Rome. Assessment results show that the maximum average concentration found un snow is 74.69 MPsL^-1^ of melted snow. The results of the characterization show that the most common MPs form was microfibers having dimensions <5mm. The largest MPs were characterized by Fourier Transformed IR (FTIR) analysis, among the polymer materials, the most abundant were polyamide (PA, 25.93%), polyethylene terephthalate (PET, 22,23%), polyester (PES, 17.28%) polypropylene (PP, 8.64%) and polyethylene (PE, 7.40%). The presence of PTFE fibers and a fragment of ABS, typical materials of technical clothing, shows that MPs contamination is also due to recreational activities as well as atmospheric deposits. The Carbonyl Index values indicated that some MPs (PE, PP) are degraded.

## Introduction

Most polymer materials spread in the environment are highly resistant to degradation therefore their complete mineralization takes hundreds of years during which degradation, mainly by photo-oxidation processes, produces microplastics and subsequently nanoplastics (Pietrelli 2024). Microplastics (MPs), particles having size in the range 0.001 – 5 mm and shapes including films, fibers, fragments and spheres, are categorized in two types based on their origin: primary, intentionally produced as small size, especially as small spheres (e.g for cosmetics), and secondary, produced following the degradation and fragmentation of macroplastics. According to the most recent scientific research it has emerged that MPs practically are ubiquitous and represent an emerging pollutants on a global scale; they can be transported by wind for long distances from the urban and industrial point sources and be later deposited, by rain or snow, in everywhere including the most remote areas, aquatic or terrestrial ecosystems (Allen et al 2019, Napper et al 2020, Gonzales-Pleiter et al 2021a, Padha et al 2022,). Even at remote sites such as Antarctic 22.5 ± 4.0 and 47.2 ± 8.4 particles per liter of melted snow have been found (Aves et al 2022) while in the snow sampled in the Everest (8440 m a.s.l.) 30±11 (94.6% of fibers) have been found (Napper et al 2020). MPs have even been found on the planetary boundaries (Gonzales-Pleiter et al 2021b). The MPs concentration can be expected to be greatest where the anthropogenic impact is relevant such as in industrial or urbanized areas: e.g. the atmospheric fallout analysed in Paris and in Hamburg shows MPs concentration till 355 and 275 pcs/m^2^/day respectively (Dris et al 2015, Dris et al 2016, Klein & Fischer 2019). Correlation between anthropic pressure and source of MPs has been observed (Crosta et al 2022), moreover MPs abundance in the remote places is correlated to their size: as fragments size decrease MPs presence increase (Hale et al 2020, Allen et al 2019, Cabrera et al 2020). Recently research on the role of MPs in the environment become a significant topic; for example, can contribute, accelerating, to the melting of snow (Ming & Wang 2021).

Up to now, most of the MPs monitoring studies have focused on their diffusion in freshwater and marine environments while, concerning high mountain ecosystems characterized by the presence of snow for long periods, the number of the published paper is relatively small (Villanova-Solano 2023). The high mountain ecosystems can be considered key areas for biodiversity, they act as water reservoirs and they are considered for many communities as cultural and religious icons (e.g. sites of “revelation”, “inspiration” and sources of springs waters and rivers). High mountains are therefore popular places and are subject to the diffusion of various pollutants such as microfibres, especially those deriving from recreational activity and, therefore, from technical equipment. At higher altitudes severe environmental conditions generally prevail and a treeless alpine vegetation is maintained. In wintertime temperatures may remain below freezing day and night permitting snow accumulation trapping micropollutants, such as microplastics, for long time therefore the snow-matrix can be considered an excellent resource to monitor the phenomenon of the MPs diffusion, especially if the sampling is done after each meteoric event: it is simple to sample and manage in the laboratory for MPs characterization.

The knowledge concerning MPs abundance and diffusion in the environment is still unclear, fragmented and probably underestimated. The main objective of this work was to obtain data regarding the diffusion of MPs in snow samples collected from the Central Apennines mountains; in addition to the abundance data, the characterization in terms of polymer identification, shape and color has been done. To the best of our knowledge this work constitutes the first article dealing with the determination of MPs in snow samples taken from these mountains that cross Italy and, in particular, from Terminillo mountain (Latium – Italy).

## Material and methods

### Study area

Monte Terminillo is massif located close to Rieti and about 100 km northeast of Rome and has a highest altitude of 2216 m a.s.l., it is surrounded by several minor ridges gently sloping towards to Thyrrenian coasts. Terminillo Mt is a typical Apennine massif in terms of morphology, fauna and vegetation and is also an active ski-resort. Mountain district that hosts typical and well-structured communities of the Apennine region with its own individuality due to some relict species and habitats. Presence of uncommon mountain habitats in the central Apennines and of many endemism can be found. According to the Habitat Directive 92/43/EEC, Terminillo mountain is a European Site of Community Importance (Monti Reatini, IT6020005, Vallone Rio Fuggio, IT 6020006, Gruppo Monte Terminillo, IT6020007, Bosco della Vallonina, IT6020009) with many habitat and species types included in the Annex I and II of the Directive such as 4060 (Alpine and Boreal heaths), 5130 (*Juniperus communis* formations on heaths or calcareous grasslands), 9180 (*Tilio-Acerion* forests of slopes, screes and ravines), 6210, (Semi-natural dry grasslands and scrubland facies on calcareous substrates, *Festuco-Brometalia* and important orchid sites) and wolf, *Canis lupus*, brown bear, *Ursus arctos*.

Concerning the sampling area, two of six sites (summit of Mt Terminillo at 2216 m a.s.l. and Rifugio Rinaldi at 2108 m a.s.l.) are characterized mainly by the presence of mountaineering tourists while for the other points, it can be observed mainly snow hiking – snowshoeing and/or ski mountaineering activities. In particular on the summit of Mt Terminillo the snow is often trampled even outside the main paths. At 1620 m a.s.l. there is an active ski resort with reduced active facilities due to the scarcity of snow that can be used to practice this sport. In Table 1 the main characteristics of the sampling sites are summarized. While in Fig. 1 the sampling areas are shown.

**Table 1.** Sampling area characteristics and abundance of MPs. Anthropic impact range 1-5.

| **Table 1.** Sampling area characteristics and abundance of MPs. Anthropogenic impact range 1-5. |  |  |  |  |  |  |  |
| --- | --- | --- | --- | --- | --- | --- | --- |
| Sampling area | Anthropic impact | Elevation<br>m a.s.l. | coordinates | MPs/<br>L | Film<br>(%) | Fragm<br>(%) | Fibers<br>(%) |
| 1) Prato Cristoforo<br>6/3/2023 | ++++ | 1875 | 42.47.66.99 N<br>13.00.88.96 E | 43.57 | 1.71 | 10.26 | 88.03 |
| 2) Sella di Leonessa<br>12/3/2023 | + | 1890 | 42.47.43.8 N<br>13.00.52.0 E | 17.91 | 0 | 7.87 | 92.14 |
| 3) Mt. Terminillo<br>16/02/2023 | +++++ | 2216 | 42.47.36.9 N<br>12.99.79.0 E | 74.69 | 1.96 | 6.72 | 91.32 |
| 4) Rifugio Rinaldi<br>12/03/2023 | +++ | 2108 | 42.46.73.3 N<br>12.99.01.3 E | 29.32 | 2.22 | 10.00 | 87.78 |
| 5) Sella di vento<br>01/03/2023 | +++ | 1629 | 42.28.02.4 N<br>12.59.24.5 E | 34.92 | 1.14 | 7.95 | 90.91 |
| 6) C. Jucci<br>28/02/2023 | +++ | 1700 | 42.88863 N<br>12.992913 E | 23.71 | 0 | 6.52 | 93.48 |

**Fig. 1.**
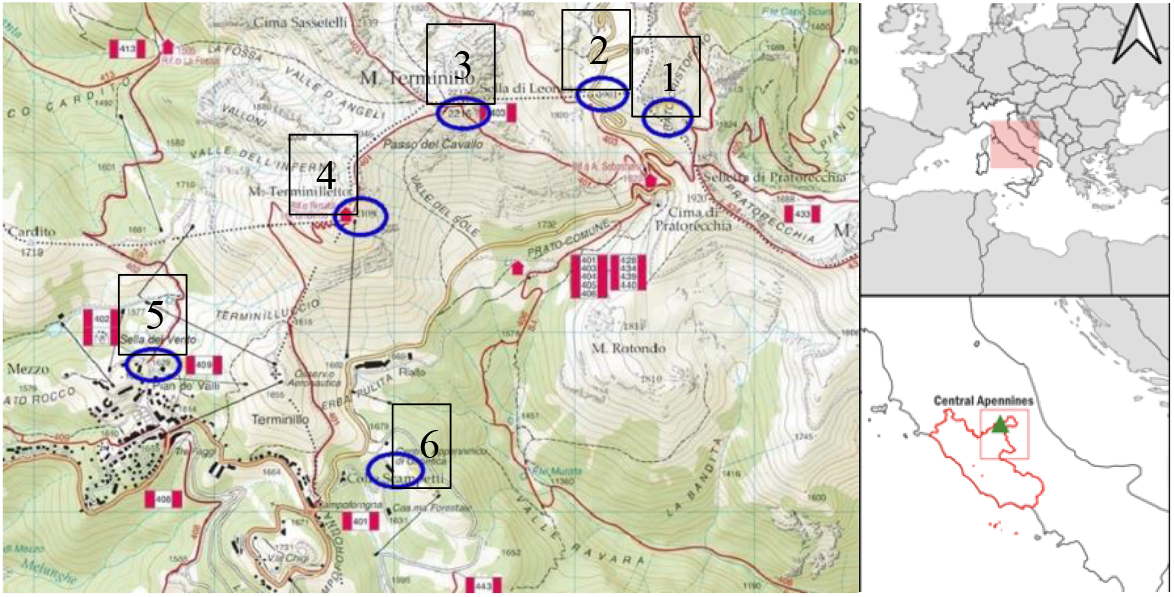
Sampling areas

### Sampling and sample treatment

Considering its simplicity, a passive sampling has been adopted (Shao et al 2022). In the period February 16^th^-March 12^th^ 2023 surface snow was sampled recovering up to 20 cm deep and quickly placed in glass bottles (1 L capacity). Each sampling site was chosen considering the absence visible contamination or movement such as footprint to avoid the human presence. To collect snow samples non-plastic sampling equipment including a white cotton overall for the operator were used, furthermore to prevent potential contamination personnel took sampling positioning themselves against the wind. For each site 5 snow samples (corners and center of a square with sides 2 m) were collected using a steel spoon (Pastorino et al 2021); totally 30 (6×5) snow samples were collected. Glass bottles were transported and stored in the chemical laboratory; snow samples were then dissolved at room temperature in a glove box to avoid external contamination. Molted snow was treated by vacuum filtration using glass microfiber filter of 0.45 μm pore size and the liquid volumes were recorded to establish the volume of each sample (Table S1, Supplementary materials). A contamination lab control sample consisting in 1L distilled water was prepared and treated together the molted snow to identify any contamination (especially from clothing) that occurred during the processing of samples. To remove the organic materials, present on the filter, Fenton’s like reaction was used (30% H_2_O_2_, Fe(II) as catalyst, pH=2-3, T≈50°C according to Aves et al (2022). The filter containing the microplastics were dried by covering the Petri dishes with a net (200 mesh) and heating in an oven at 60°C for 24 hours. A labelled glass Petri plate was used to store the filter containing microplastics in freezer until analysis. To simplify the MPs quantification and characterization, on the surface of the filter a grid with squares of 5 mm side was drawn (Fig. 2).

**Fig. 2.**
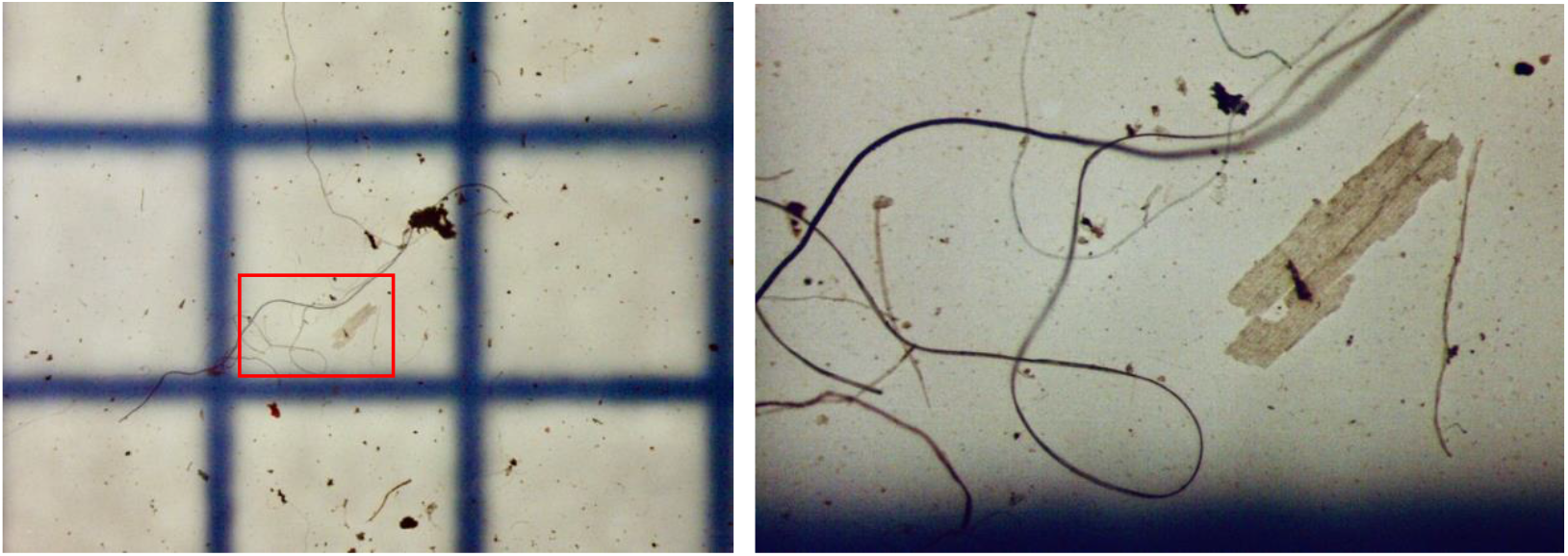
Microplastics deposited on the filters after the chemical treatment observed by optical microscopy. In the enlarged box (25x) it is possible to observe, together with various fibres, the highly degraded LDPE fragment

### Microscopic observation

After sample treatment filters were inspected using an optical microscope Leica K 200 LED. Considering the difficulty in distinguishing dark colors such as black, brown, gray etc, these colors have been grouped into a single “dark” category. To enhance the MPs visibility, the filters were illuminated by white LED light and MPs were counted in each 5×5 mm square. MPs were classified by shape, color and dimension (less or greater than 5 mm).

### Characterization by FTIR

The recovered items were classified and characterized according to Galgani et al (2013); shape (fibers, film, fragments and beads) and color. MP particles characterization refers to the principal chemical component of the polymer, therefore characterization of the largest samples was carried out with Fourier Transformed Infrared Spectroscopy (FTIR). A Thermo-Scientific Nicolette 6700 FTIR spectrophotometer was used. Due to the small size of most recovered MPs it was possible to chemically characterize by FTIR only 81 samples (10.4 %) (instrumental detection limit ≈0.3 mm).

The measurements were carried out in total reflection with the use of a single reflection ATR accessory (model Golden Gate Single Reflection ATR System). The chosen spectral range was from 4000 cm^-1^ to 650 cm^-1^, with a resolution of 4 cm^-1^ and with an accumulation number of 100 per spectrum. The polymers were identified considering a correspondence between the measured spectrum (sample) and the reference one, this latter present in the instrument database, between 75 and 95%. Following the degradation processes, new peaks appear on the IR spectrum: a peak at a region of 3300-3400 cm^-1^ (-OH, hydroxyl groups), at 1650–1800cm^-1^ (C=O, carbonyl groups, visible in ketones, carboxylic acids, esters and centered, for saturated compounds, at 1715 cm-1) at 1600-1680 cm^-1^ and at 909 cm^-1^ (C=C, carbon double bonds) and 1000-1250 cm^-1^ (C-O-C ether groups). The formation of carbonyl compound represnts a sign of oxidation of polyolefins which helps to enhance the degradation process. Therefore, using the IR spectrum, the polyethylene (PE) and polypropylene (PP) degradation was revealed using the Carbonyl Index (CI= *A*(1720) / *A*(722)) that was calculated using the absorption band at 1720 cm^−1^, stretching vibration of the carbonyl group (C=O) while the absorbance at 722 cm^−1^ is used as reference (Roy et al 2007, Pietrelli 2024). The CI is taken as the average of at least three random samples from the MPs.

### Meteorological data

The meteorological observations during the sampling period are those collected at the Colle Scampetti weather station (1730 m a.s.l. 42°27’15”N; 00°32’30” by M. Mario) and made available by the Terminillo “Carlo Jucci” Apennine Center (Rieti).

In the period from 02/03/2023 to 29/04/2023 it snowed for 14 days for a total of 57 cm of snow, while the cumulative rain was 285 mm. The minimum temperature of the period was -6.2°C and the maximum was 13.8°C, while the average relative humidity was 71%, with peaks of 98%. During the sampling period, wind speeds often exceeded 60 km/h per hour with peaks of over 80 km/h (maximum 85.2 km/h). The prevailing wind directions were NW, W.

## Results and discussion

Following the melted snow filtration totally 1651 microparticles were separated, the average value of organic material particles oxidized with the Fenton’s like reaction was 52.9% therefore the MPs present in the snow samples collected during the monitoring campaign were 777 in total (Table S1). The high quantity of microplastics present in the snow (till 74.69 MPs/L) that fell just during few days (14) is not surprising; even in samples collected in more remote areas (less visited by tourists) a high number of MPs can be found (17.91 MPs/L) (Table 1).

Regarding the type of microplastics found in the snow samples, this monitoring campaign suggest that microfibres (MFs) concentration is higher than any other type as reported in Table 1. No microbeads were found indicating the absence of cosmetic products. As expected, films and fragments were more abundant close to the sampling sites most frequented by tourists and hikers, even the quantity of larger fibers is more abundant in places where human presence is greater. In particular, the presence of microfibers longer than 5 mm would suggest that they are not the result of atmospheric fallout but that they derive from the presence of recreational activities such as skiing and hiking (technical fabris). After all, Monte Terminillo is one of the favorite destinations for the Rome inhabitants (about 3 millions of inhabitants at about 110 km away) and the presence of ABS fragments probably due to the wear of the soles of technical shoes could confirm this hypothesis (figure 1S supplement materials). In contrast, in other European mountain ranges such as the Pyrenees mountains has been found that the snowfall events are correlated with the MPs depositions and the most abundant MPs were found to be <1.2 mm (Allen et al 2019).

The histogram in Fig. 3 shows the color, size and number of MPs found in snow samples. Considering MPs as single size, dark colors were the most abundant (38.56%) followed by blue (22.87%), transparent/white (18.49%), red (14.36%) and purple (5.72%). Making a comparison with other published papers we can see that the prevailing colors are black and blue in Spain, Antarctica sites (Villanova-Solano et al 2023, Bergami et al 2023) and on Tibetan Plateau where MPs are like those found in the urban airborne (Zhang et al 2021).

**Fig. 3.**
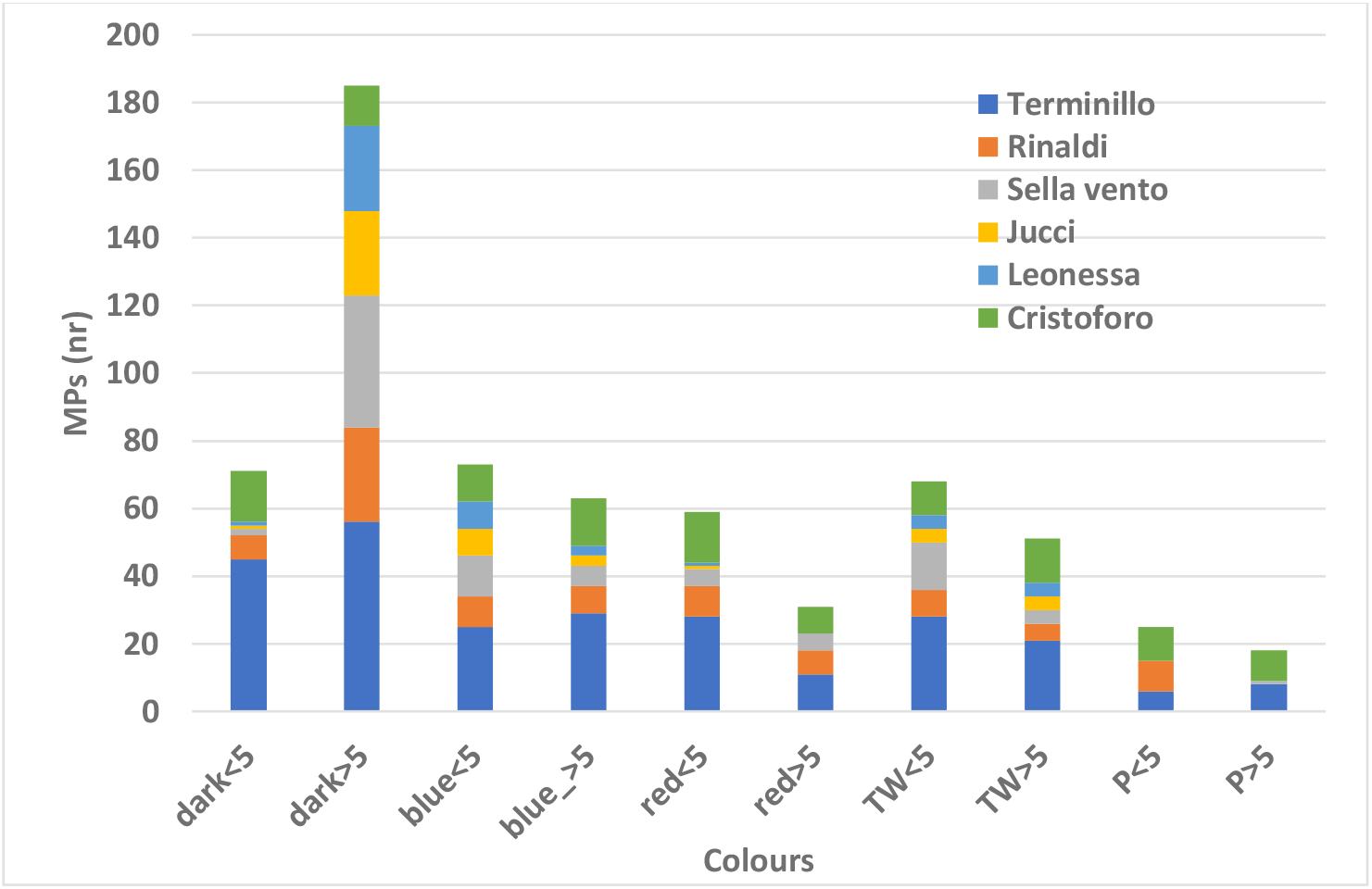
Colors distribution of the total fibers found in each sampling point according to dimension (> or < 5mm). Dark= black, brown and grey, TW=transparence and white color, P=purple

Table 2 shows a comparison with several papers reporting the abundance and the characterization of MPs in snow in high altitude and close to large cities. They indicate that microfibers are the shape most frequently observed in the snow samples. Data regarding microplastics concentration in terms of MPsL^-1^ seems to be of the same order of magnitude as those obtained by Yu et al in Mongolia (2023), in Russia by Chernykh et al (2022) and Villanova-Solano at 2150 m a.s.l. at Tenerife (2023) close to urban and industrial areas.

**Table 2.** Worldwide MPs abundance in the snow. Polymers are listed based on their abundance

|  |  |  |  |  |  |  |
| --- | --- | --- | --- | --- | --- | --- |
| Pathways | Where | Altitude<br>m a.s.l. | Abund.<br>MPsL<sup>-1</sup> | Shapes | Polymers | Reference |
|  | W.Alps Italy | >2500 | 4.9±2.5 | Fibers<br>fragments | PE, PET, PES, PC, PP,<br>PU | Parolini et al<br>2021 |
|  | Forni Ice<br>Alps Italy | 2580 | 70±32.9<br>MPsKg<sup>-1</sup> | Fibers<br>fragments | PES, PA, PE, PP | Ambrosini et al<br>2019 |
| B | Antisana ice<br>Equador | 5753 | 101.2 | Fibers, Film<br>fragments<br>sphere | ? | Cabrera et al<br>2020 |
| B | Sonnblick<br>obs.Austria | 3106 | 46.5 mg/L | Sphere, fibers | PET, PP, PS, PE | Materic et al 2021 |
| A | Carnic Alps<br>Italy | 1872 | 0.11±0.19 | fragments | PET | Pastorino et al<br>2021 |
| A, B | Alaknanda<br>India | 1524 |  | Fragments<br>fibers | Fragments<br>fibres | Stefansson et al<br>2021<br>Chauhan et al<br>2021 |
| A | M.Everest<br>Nepal | 8440 | 30±11 | Fibers<br>fragments | PES, Acrylic, Nylon,<br>PP | Napper et al 2020 |
| C | Barnaul<br>Russia | 180 | 27-595 | Foam,<br>fragments,<br>film, fibers |  | Chernykh et al<br>2022 |
|  | Swiss Alps | 2019 | 9800±6900 | fibers | PE, PP, PS, PC, PA,<br>PVC, Cell., PES,<br>Nylon, Rubbers, EVA | Bergmann et al<br>2019 |
|  | Tibetan<br>plateau | 4460-5600 |  | Fibers<br>fragments | PA, PE, rubbers, PET,<br>PS, PC, PMMA | Zhang et al 2021 |
|  | Tabriz Iran | 1350 | 20 | Fibers, Film fragments sphere | Nylon, Cell., PP, PES, PET, EVA, PS, PVC, PE | Abbasi et al 2022 |
|  | Ross Island |  | 29.4±4.7 | Fibers, Film fragments | PET, PMMA, PVC, PA, PE, cell | Aves et al 2022 |
| A | Tenerife Spain | 2150<br>3200 | 51±72<br>167±104 | Fibers, Film fragments | Cell., PES, Acrylic, Nylon, Azlon, PP | Villanova-Solano et al 2023 |
| C | Inner Mogolian plateau China |  | 68±10<br>199±22 | Fibers (>80%) Fregments foam | PE, PA, PES, PS, PVC, PET | Yu et al 2023 |
| C | Hokkaido Japan | 5-1143 | 150<br>4200 | Fragments (97%) | PE, rubber, PA, PVC, PMMA, rayon, PU, EVA | Ohno & Lizuka 2023 |
| C | Ankara | 5-938 | 59.66 | Fibers, Film fragments | PE, PS, PP, PET, PVC, Nylon | Babaei et al 2023 |
| A, B | 2217 |  | 43.86<br>163.18 |  | PA, PP, PE, PET, PES, Cell., PTFE, SB, PA | This study |
| A= Anthropogenic activity, B=atmospheric deposition, C=close to industrial activity or city |  |  |  |  |  |  |

Considering anthropogenic materials, totally we detected 9 different polymer types and the only site where all the polymers have been found is the summit of Mount Terminillo. Fig. 4 shows the result of the MPs characterization; the most abundant were polyamide (PA, 25.93%), polyethylene terephthalate (PET, 22,23%), polyester (PES, 17.28%) polypropylene (PP, 8.64%), polyethylene (PE, 7.40%), cellophane, polytetrafluoethylene (PTFE), Nylon and styrene-butadiene copolymer. Most of the plastic items found in the snow were in the form of fiber probably from textile (Table 1) in accordance with what has been reported by numerous other authors as shown in Table 2. We observed differences in size (<5mm>) when comparing plastic items in the collected samples (T=21, P<0.01, Wilcoxon two-sample paired test). In particular, on the summit of Mount Terminillo more microplastics were found and furthermore, unlike the other sampling points, more fibers with dimensions greater than 5 mm were found. This is probably because this site is reached by many hikers throughout the year and therefore the quantity of smallest MPs transported by the wind is lower in percentage.

**Fig. 4.**
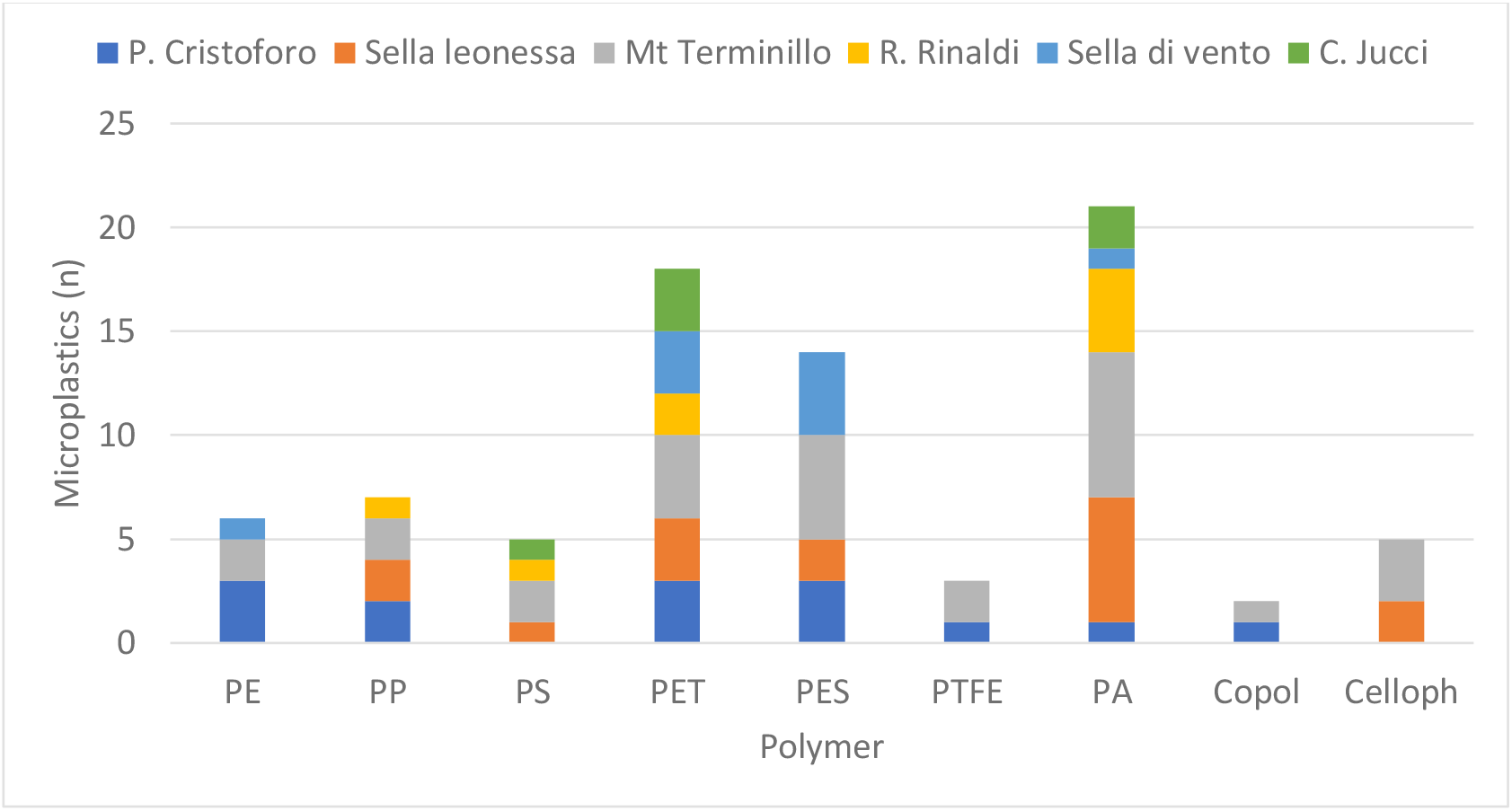
Polymer materials found in snow samples.

The diffusion of polymers such as PP and PE is not surprising since within the fossil-based polymers, polyethylene (HDPE+LDPE) and polypropylene (PP) are the most produced synthetic polymers with respectively 26.3 and 18.9 % of the total polymer materials produced (PlasticEurope 2023). While the presence of other polymers in fiber form such as PTFE, PA and polyesters can be attributed to the strong presence of mountaineering, snow hiking, snow shoeing and ski mountaineering who frequent those mountains during each season. Many fibers found on the top of the mountain are due to the fraying of the flags placed on the cross that “welcomes” hikers (Figure S1 of supplementary materials) Among the various polymeric materials, the presence of PTFE (Fig. S2 of supplementary materials), the constituent of Gore-tex^®^, a patented type and highly popular waterproof material, it was found in the Antarctic snow (Aves et al 2022). Some fragments of elastomers (Styrene-butadiene, isoprene) were also found in the snow samples probably originating from the soles of the boots (Fig. S3 and S4 Supplementary materials).

The characterization of the collected samples clearly shows that some polymer materials, in particular PE and PP are degraded due to long permanence in the environment and subjected to the action of photodegradation processes (Fig. 5 and 6). The structural variations observed through the FTIR spectra of the degraded PE and PP demonstrated by several peaks: in the region of 3300-3400 cm^-1^ attributable to hydroxyl groups (-OH) in the range of 1600-1800 cm^-1^ attributable to carbonyl groups (C=O) and in the range 1000-1250 cm^-1^ characterized by the presence of the ether groups (C-O-C). The presence of these oxygen-containing groups is the consequence of photodegradation reactions due to simultaneous exposure to the sun and air. Further evidence of degradation can be obtained thanks to the carbonyl indexes that were in the range 0.3-0.4 and 0.2-0.5 respectively for PE and PP, these values are comparable to that found by other authors (Tocháček & Vrátníčková 2014, Pietrelli 2024).

**Fig. 5.**
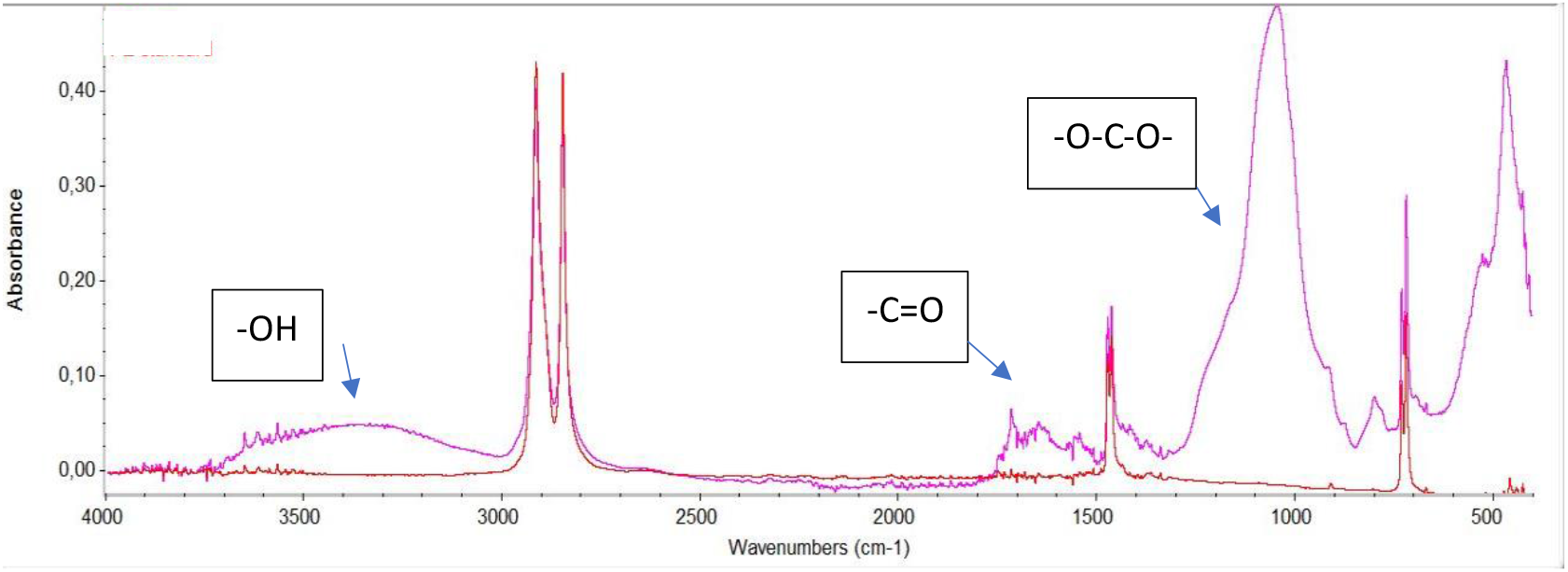
IR spectra of standard LDPE (red color) and PE film fragment collected in snow (purple color)

**Fig. 6.**
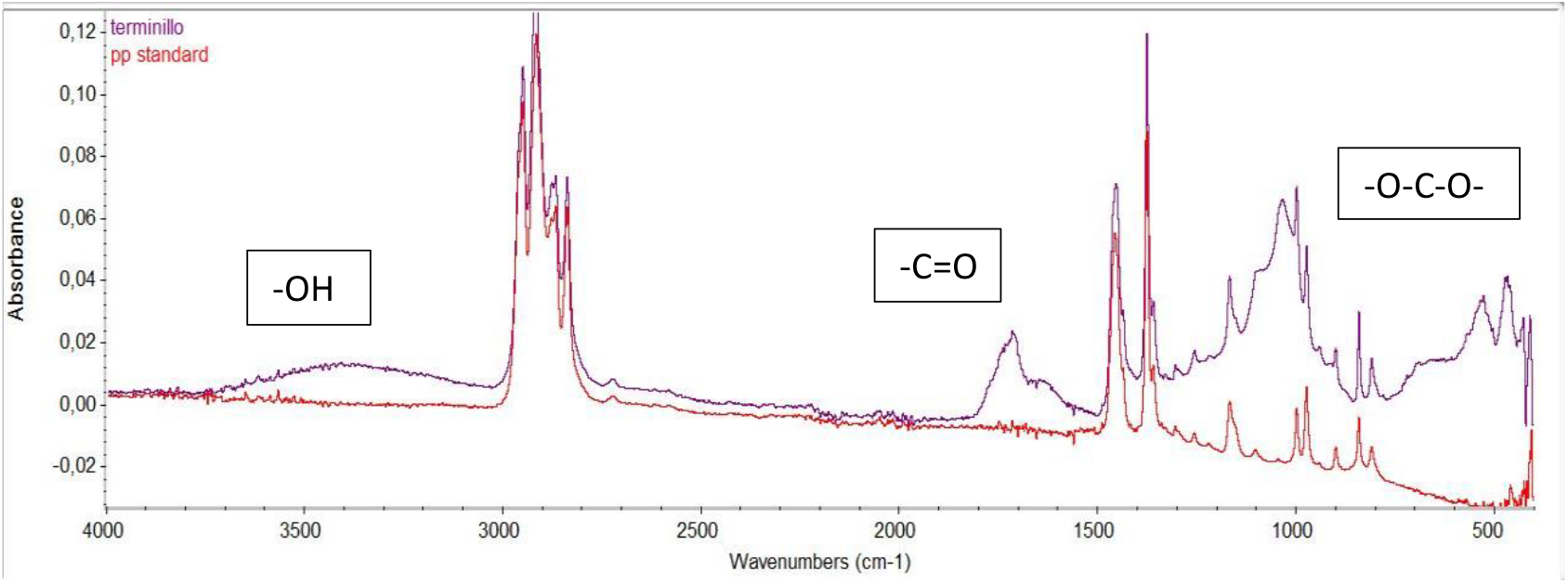
IR spectra of standard PP (red color) and PP film fragment collected in snow (blu color)

## Conclusions

The analysis of snow sampled from the Terminillo massif confirms the extent of the phenomenon related to the spread of microplastics in the environment indicating the severity of the atmosphere as a source of airborne MPs issue. To this phenomenon it is necessary to add the contribution due to significant recreational activities such as hiking, alpine skiing. It is also important to consider that in addition to the presence of polymeric materials microplastics can have harmful substances as original constituents such as filler, plasticizers, flame retardants and other substances adsorbed on to their surfaces such as heavy metals, bacteria, persistent organic pollutants such as pesticides and hydrocarbons (Chauhan et al 2021).

MPs in snow have fallen out, 17.91-74.69 MPsL^-1^ among which dark and blue microfibers having size less than 5 mm were particularly abundant, suggesting that this pollution could be coming from long distance sources. The presence of larger MPs especially where the tourist influx is greater concerns the presence of local sources (recreational activities). The amount and quality of data obtained collecting sample of snow can be considered, therefore, an indicator of airborne MPs especially because it is easy to sample. Moreover, considering that each recreational activity such as ski mountaineering, hiking or simple sightseeing is associated with appropriate technical clothing, from the analysis of the MPs found one could also get an idea of the prevalent activities. Our results confirmed that local contamination can represent a relevant source of MPs in mountain ecosystems due to the high anthropic pressure, while long-range transport can be the main source on site where the human presence is absent or limited. Considering the impact of MPs on the environment and in particular, on the soil once the snow melt, future research should be conducted on the high mountain ecosystems following the fate of MPs. To understand the environmental fate of MPs in high-mountain ecosystems an increase of monitoring project should be done at best using a citizen science approach. Considering that it is impossible to remove PMs from environment, solutions for plastic pollution reduction it is necessary to apply many actions such as re-use, recycling, reduction (the 3-R approach) accompanied, however, by an extensive awareness campaign aimed at citizens.

## Authors contributions

All authors contributed to the study conception and design. Study design L.P., P.M., B.S., Sampling I.M., Material preparation, data collection and analysis were performed by P.M., M.S., M.B. and L.P. The first draft of the manuscript was written by L.P., and P.M. All authors commented on previous versions of the manuscript. All authors read and approved the final manuscript.

## Data availability

The authors declare that the data supporting the findings of this study are available within the paper and its Supplementary Information files. Should any raw data files be needed in another format, they are available from the corresponding author (L.P.) upon reasonable request.

## Ethical Approval

Not applicable. *Consent to Partecipate* Not applicable.

## Consent to Publish

All the authors are willing to publish this paper in ESPR.

