## supplemental material for "MICROPLASTICS IN THE MOUNT TERMINILLO SNOW’S (RIETI, ITALY)"

Table S1: Total particles (mps) and Microplastics (MPs) found in the snow samples

| Sample | Volume (mL) | mps | MPs | sample | Volume (mL) | mps | MPs |
| --- | --- | --- | --- | --- | --- | --- | --- |
| Terminillo (2216 m a.s.l.) | | | | Centro Jucci (1700 m a.s.l.) | | | |
| W | 800 | 207 | 166 | W | 220 | 30 | 12 |
| North | 980 | 51 | 18 | North | 290 | 12 | 7 |
| Est | 1000 | 94 | 21 | Est | 290 | 28 | 9 |
| Sud | 1000 | 165 | 57 | Sud | 810 | 26 | 12 |
| Center | 1000 | 263 | 85 | Center | 330 | 6 | 6 |
| **Total** | **4780** | **780** | **347** | **Total** | **1940** | **102** | **46** |
| Rifugio Rinaldi (2108 m a.s.l.) | | | | Prato Cristofaro (1875 m a.s.l.) | | | |
| W | 1000 | 55 | 34 | W | 815 | 58 | 32 |
| North | 520 | 71 | 27 | North | 340 | 40 | 28 |
| Est | 550 | 17 | 11 | Est | 240 | 33 | 18 |
| Sud | 1000 | 64 | 18 | Sud | 640 | 25 | 13 |
| Center | No | 0 | 0 | Center | 650 | 32 | 26 |
| **Total** | **3070** | **207** | **90** |  | **2685** | **201** | **117** |
| Sella del vento (1890 m a.s.l.) | | |  | Sella Leonessa (1875 m a.s.l.) | | | |
| W | 660 | 32 | 14 | W | 1260 | 44 | 18 |
| North | 230 | 36 | 20 | North | 620 | 38 | 21 |
| Est | 580 | 22 | 13 | Est | 660 | 26 | 9 |
| Sud | 600 | 28 | 21 | Sud | 1380 | 65 | 29 |
| Center | 450 | 25 | 20 | Center | 1050 | 45 | 12 |
| **Total** | **2520** | **143** | **88** | **Total** | **4970** | **218** | **89** |

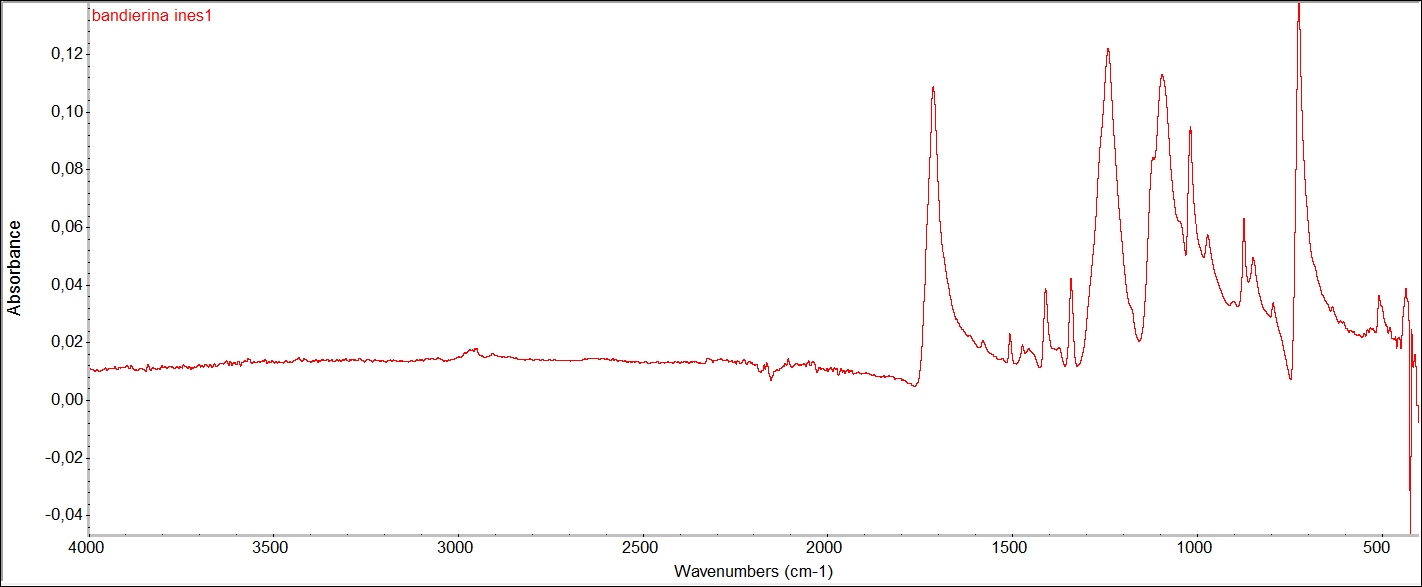

**Fig. S1**. The FTIR spectra of the polyester fiber form flag

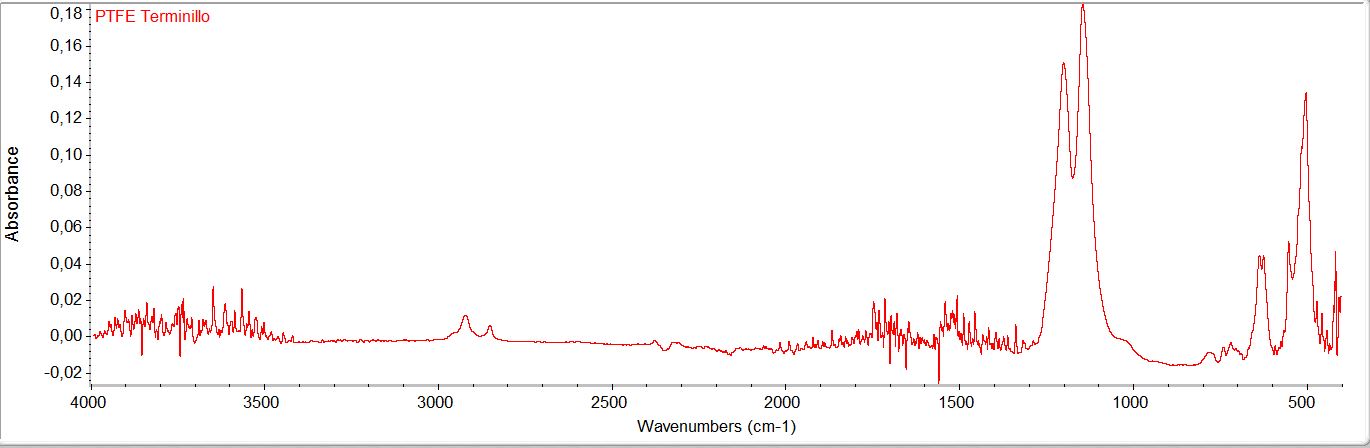

**Fig. S2**. The FTIR spectra of the PTFE fiber

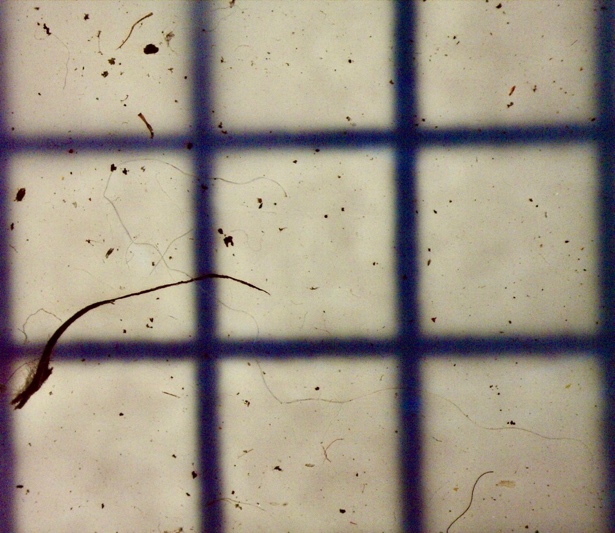

**Fig. S3**. The ABS fragment found in snow

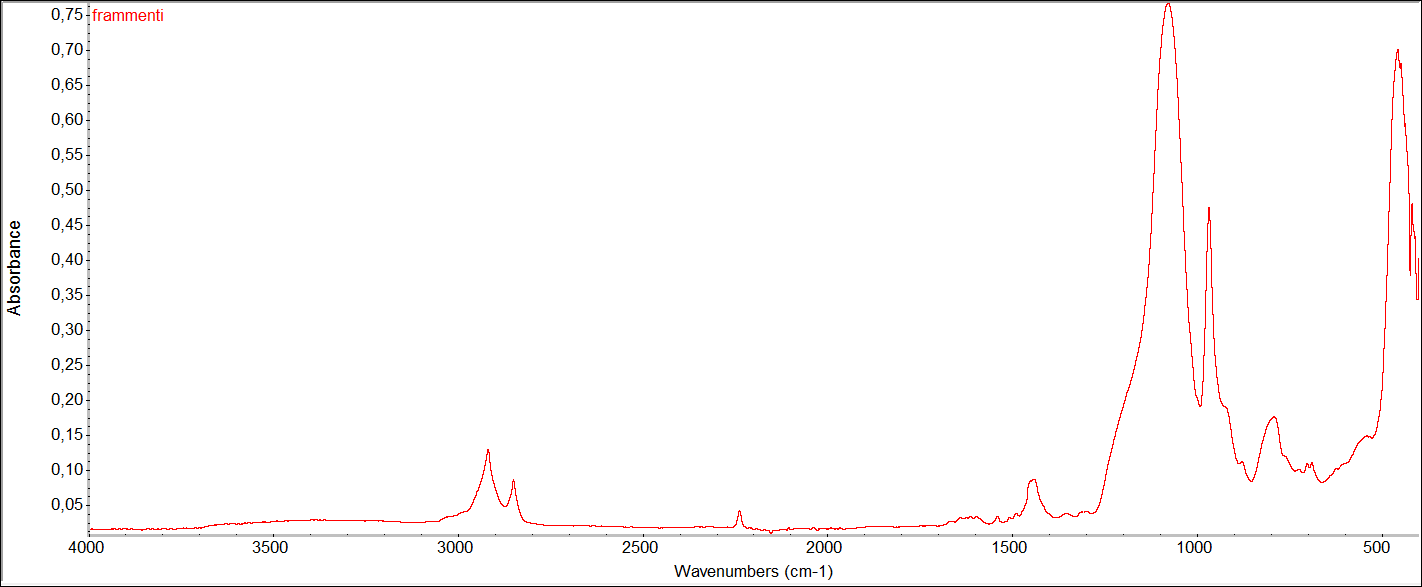

**Fig. S4**. The FTIR spectra of the ABS fragment
